# A functional Red List Index for monitoring trends in functional diversity

**DOI:** 10.1101/2025.10.30.685639

**Authors:** Kerry Stewart, Joseph A. Tobias, Stuart H. M. Butchart, Chris Venditti, Carlos P. Carmona, Chris Clements, Manuela Gonzalez-Suarez

## Abstract

Functional diversity supports ecosystem function and resilience but is in decline due to ongoing extinctions. Despite this, no functional diversity metrics are included in the set of indicators adopted by governments to measure progress towards the Global Biodiversity Framework. Monitoring changes in functional diversity is essential to understand differences between taxonomic and functional diversity trends. The Red List Index (RLI) is a widely used metric in conservation policy, and tracks trends in aggregate extinction risk across sets of species over time. We present a functional Red List Index (fRLI), which weights species by their morphological uniqueness at either local or global scales, to indicate the risk that future extinctions could pose to functional diversity. We applied the fRLI on birds and found higher aggregate extinction risk for both local and global fRLI compared with the standard RLI. Species with higher local uniqueness were more likely to be at higher risk of extinction while we found no difference for globally unique species. Globally or locally unique species were not more likely to show escalation in extinction risk between 1988 and 2024 than less unique species; however, locally unique birds had a lower proportion of their resident range covered by site-based conservation measures, leaving them vulnerable to future decline. The fRLI provides a tool to monitor the impact of extinction risk on functional diversity and could be applied to other taxonomic groups with comprehensive or representative coverage of Red List assessments and trait data.

## Introduction

Maintenance of ecosystem functioning and resilience is essential for human societies, with implications for water availability, food security, and human health^1^. Even so, over 46,000 species are known to be threatened with extinction and extrapolations suggest that as many as 1 million species may be threatened with extinction^2^, as human activity continues to reduce the viability of species populations^3^. In response, governments adopted the Kunming-Montreal Global Biodiversity Framework in 2022, to halt and reverse biodiversity loss by 2030 and put nature on a path to recovery for the benefit of people and planet^4^. To monitor progress towards this goal, the Parties to the Convention on Biological Diversity adopted a monitoring framework, with headline, component and complementary indicators^5^. However, there are no metrics of functional diversity (describing the diversity of traits in an ecosystem that influence an organism’s niche^6^) in the framework. There is wider concern that metrics developed and used for monitoring the impacts of species conservation may provide limited insight into the status and conservation of key ecological functions and processes^7–9^.

A possible solution is to integrate functional traits into current species-focused indicators so that they are more informative about the functional implications of biodiversity decline. Functional diversity often provides a better descriptor of ecosystem functioning than species richness^10,11^, so integrating functional traits into biodiversity metrics could provide a better understanding of the impact of changes in extinction risk on the status of key ecosystem processes^10^ and ecosystem resilience than species-based metrics alone^12,13^ (although see^14^). The Red List Index (RLI)^15–17^ is a well-established species-based metric, which uses data from the IUCN Red List: a standardised method for classifying species’ extinction risk, informing conservation efforts from local to global scales^18^. The RLI has been central to monitoring the state of the biodiversity crisis over the last two decades^17^, and it is widely used in policy and practice, including as a headline indicator for Goal A and Target 4 of the Kunming-Montreal Global Biodiversity Framework^4^, and for tracking progress towards target 15.5 (to halt the loss of biodiversity) of the United Nations Sustainable Development Goals^19^. In the global RLI, all species within a taxonomic group are treated equally and given a weighting according to their extinction risk category^20^. Here we propose a functional Red List Index (fRLI) that weights species according to the uniqueness of their functional traits to provide a measure of the impact of species’ declines on functional diversity, helping to fill the gap in metric availability for monitoring functional diversity change for conservation^21^.

Projected extinctions pose a disproportionate risk to functional diversity^22^, owing to the fact that functionally unique species are at greater risk of extinction than less functionally unique species^23,24^. Species that are functionally unique are important for maintaining future delivery of ecosystem services (maintaining “option value”), and globally distinct species are more likely to be utilised by humans for food, medicine, materials and other economic and cultural purposes^25^. Importantly, species that are morphologically similar to many other species within the global avian assemblage may still be unique within their habitat or range and provide a unique contribution to ecosystem function, so any attempts to integrate functional traits into biodiversity monitoring must be scale-explicit. While unique species are currently more threatened with extinction than less unique species^23,24^, we have limited understanding of whether unique species are more or less likely to be moving closer to extinction over time. Functional diversity will be better conserved if unique species are less likely to be experiencing escalations in extinction risk or are more likely to be experiencing recovery, while it could be substantially eroded if unique species are more likely to have elevated extinction risk or less likely to be recovering (figure 1).

**Figure 1.**
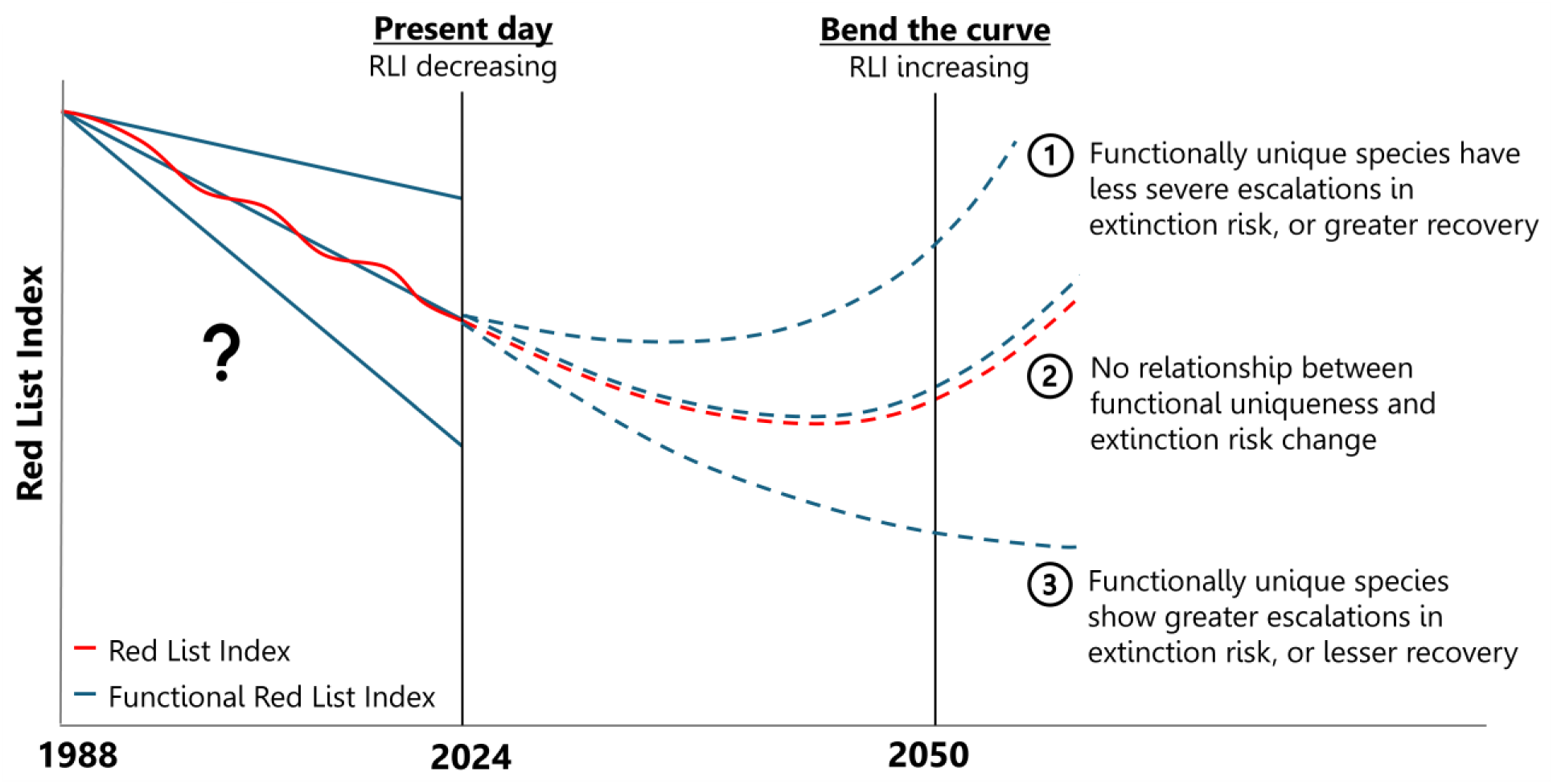
Potential past and future trajectories of Red List Indices. Both the existing RLI and our proposed functional Red List Index (fRLI) are shown under three scenarios. RLIs show declines across all major taxonomic groups^26^. To bend the curve on biodiversity loss, species extinctions must be halted and recovery promoted^4^, which would drive an increase in the RLI^20^ (indicating a reduction in average extinction risk across species). Scenario 1: if functionally unique species have less severe escalations in extinction risk or show greater recovery, the fRLI may begin increasing before the RLI. Scenario 2: the fRLI may follow the same trajectory as the RLI if changes in extinction risk are not biased with respect to species traits. Scenario 3: if functionally unique species experience greater declines in extinction risk, or lesser recovery than less unique species, the fRLI may continue to decline even if the RLI begins to increase.

Our proposed functional Red List Index (fRLI) monitors aggregate change in extinction risk over time, explicitly considering species’ functional uniqueness. This metric complements the existing RLI and similarly uses information on genuine changes in extinction risk^26^, but additionally uses data on species’ traits to identify the local and global uniqueness of species. We apply this metric to birds, the best-known class of organisms with well-documented trends in extinction risk^17^ and detailed data available on their traits^27^, to explore whether changes in extinction risk have had a greater, or lesser, impact on functional diversity than species richness (i.e. comparing fRLI with RLI). To better evaluate factors influencing differences between the two indices, we tested whether globally and locally unique birds had higher or lower extinction risk in 2024, were more or less likely to have undergone genuine reductions or increases in extinction risk since 1988 (when birds were first assessed for the IUCN Red List), and whether they were more or less likely to be covered by site-based conservation measures. Globally or locally unique species may be more likely to be covered by site-based conservation measures if they receive greater human attention due to their unusual morphology^28^, while they may have lower coverage if they occupy rare niches so benefit less from conservation measures in place to protect other species^29^. In this study we aim to (i) estimate the impact of changing extinction risk on functional diversity and associated ecosystem function, and (ii) assess the extent to which existing conservation approaches address the multifaceted nature of biodiversity.

## Results and Discussion

### Functional uniqueness

We used principal component analysis to summarise variation in eleven morphological traits, obtained from a published dataset^27^, which describe avian ecological niches through their association with diet^30^, dispersal^31^ and habitat^32^ (see Methods). Since four independent axes of morphological trait variation are the minimum required to predict avian trophic niches^30^ we selected the first four principal components that described 93.7% of variation in body, wing, tail, beak, and tarsus morphology. The first principal component was primarily associated with body size; the second was associated with wing morphology; the third described variation in beak length and to a lesser extent tail length; and the fourth described beak width and depth (and to some extent tarsus length, Supplementary table 1).

We calculated functional uniqueness of species using trait probability densities^33^ following the method of Stewart et al.^34^ (figure 2). This metric describes the probability that functional richness is lost when a species goes extinct. Species may be unique at a global scale (very different morphologically from other extant species) or unique at a local scale (very different morphologically from species with which they co-occur, even if potentially similar to other species living elsewhere). To capture both scales, we defined *global uniqueness* as the functional uniqueness of species within the global avian assemblage, and *local uniqueness* as the functional uniqueness within local terrestrial avian assemblages. When calculating global uniqueness, the community probability distribution was built with all 11,038 species for which we had measured or inferred trait data (8.2% of species had inferred data for at least one trait). When calculating local uniqueness, a community probability distribution was built for each 110 x 110km terrestrial grid cell across the world, using species distribution data from area of habitat maps from BirdLife International^35^.

**Figure 2.**
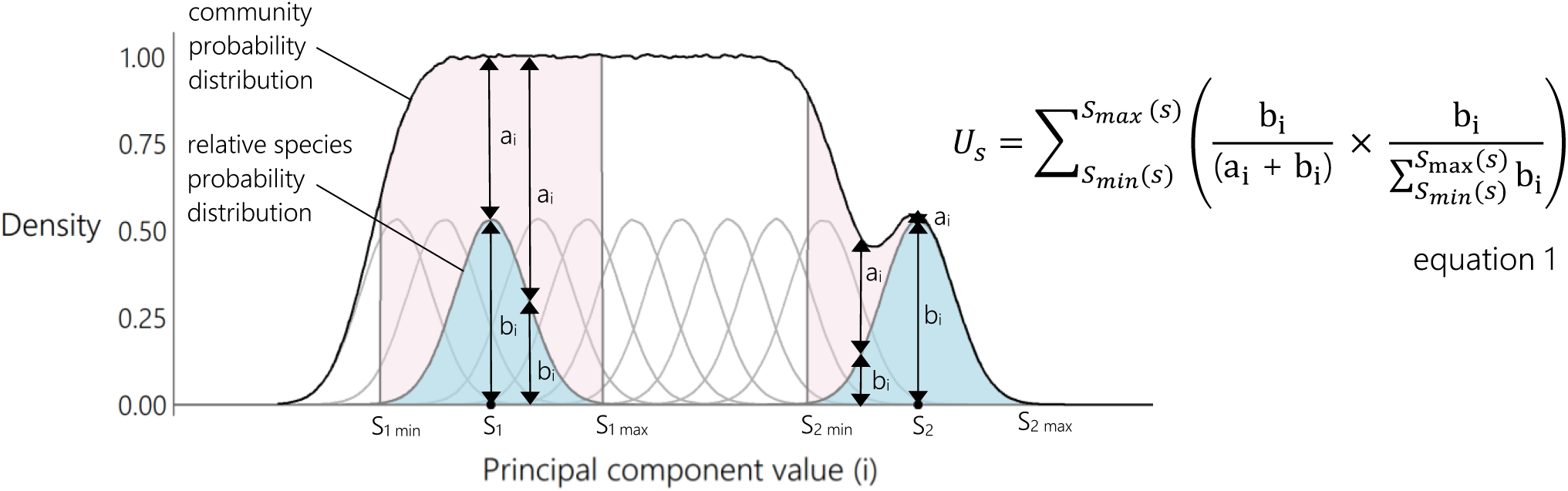
Method for calculating functional uniqueness. Individuals from species S_1_ are likely to occur between S_1min_ and S_1max_, where the relative species probability distribution is greater than 0 (species probability distributions are truncated to include 0.95 of the total species probability distributions). Between S_1min_ and S_1max_, the relative species probability distribution of S_1_ overlaps with six other relative species probability distributions, so the density of the S_1_ relative species probability distribution makes up a small to moderate proportion of the density of the community probability distribution (b_i_ similar magnitude to a_i_ at the peak of the S_1_ relative probability distribution and smaller than a_1_ at all other parts of the relative species probability distribution). On the contrary, the S_2_ relative species probability distribution overlaps with three other relative species probability distributions and makes up a large proportion of the community probability distribution between S_2min_ and S_2max_. In this case, species S_2_ has higher uniqueness than S_1_ because functional richness loss would be more likely if species S_2_ went extinct than if species S_1_ went extinct, owing to the low overlap of the S_2_ relative species probability distribution with other relative species probability distributions.

Based on trait probability densities, 369 species (3.3% of all study species) were found to be morphologically unique – that is the species probability distributions of these species showed no overlap with the species probability distribution of any other birds in the defined gridded four-dimensional trait space (figure 3). Species probability distributions were multivariate gaussian distributions with means obtained from principal component values derived from functional trait data^27^ and standard deviations that were estimated using a bandwidth selector^36^. Sensitivity analyses revealed that our results were robust to deviation in estimated standard deviations of traits (Supplementary figure 1 and 2).

**Figure 3.**
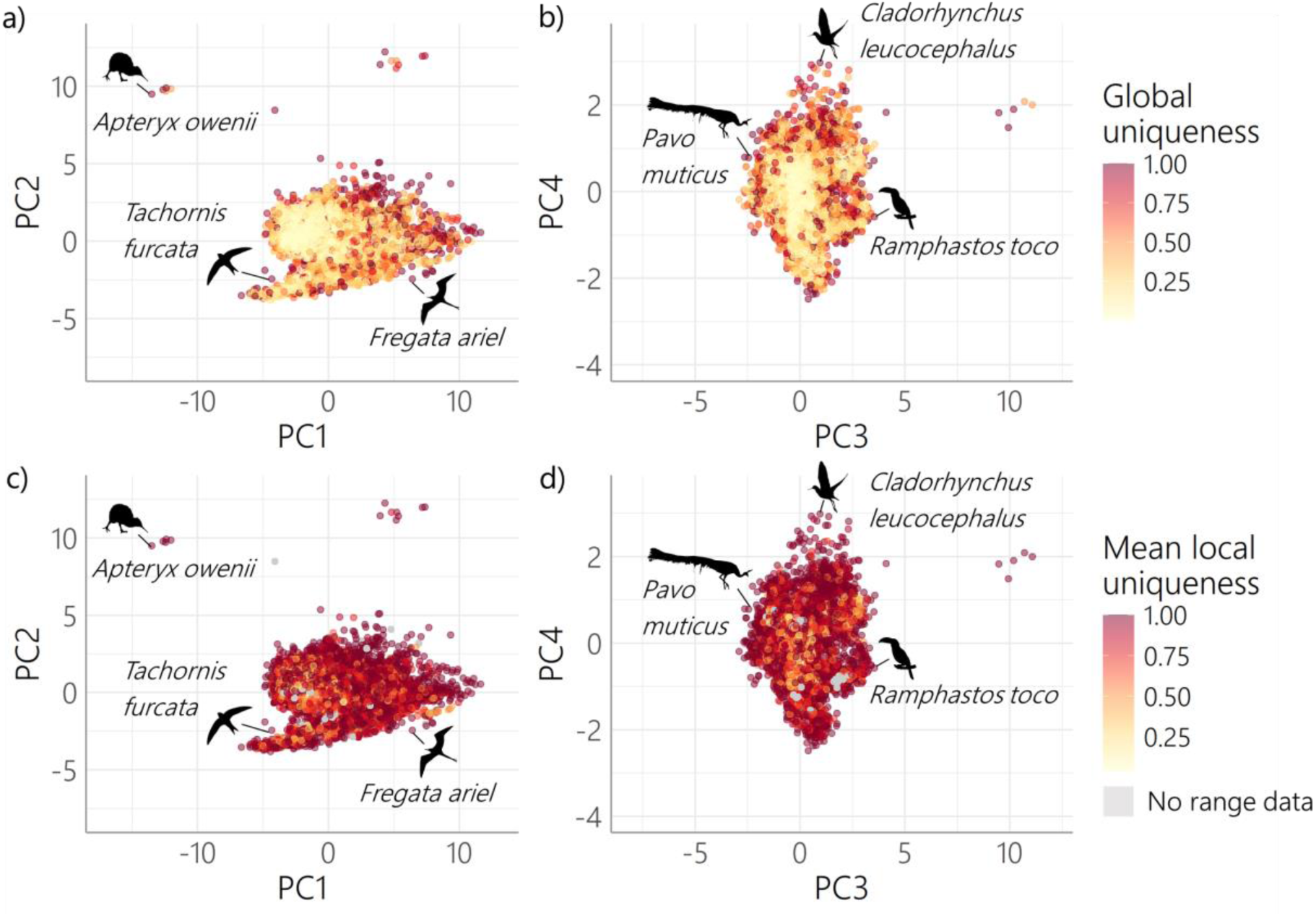
The distribution of global uniqueness and mean local uniqueness among the world’s birds. Global uniqueness calculated across four principal components shown in relation to **a,** PC1 and PC2 and **c**, PC3 and PC4, and mean local uniqueness shown in relation to **b,** PC1 and PC2, and **d,** PC3 and PC4 (n=11,038 species for all panels). Six species which have a global and mean local uniqueness of 1 have been highlighted. For 179 species it was not possible to calculate local uniqueness (grey points in panels c and d, no range data) because they had no terrestrial range (145 species, including those restricted to small islands or ice shelves and sea ice), or no known remaining extant range (34 species). All silhouettes from <u>Phylopic</u>. Panel a and c left to right: *Apteryx owenii* (*Apteryx* by Ferran Sayol, CC0 1.0), *Tachornis furcata* (*Apus apus* by Ferran Sayol, CC0 1.0), *Fregata ariel* (*Fregata magnificens* by Emily Willoughby, <u>CC BY-SA 3.0</u>). Panel b and d left to right: *Pavo muticus* (*Pavo muticus* by Holly Marshall, CC0 1.0), *Cladorhynchus leucocephalus* (*Cladorhynchus leucocephalus,* by T. Michael Keesey from a photograph by Ed Dunens, CC0 by 4.0), Ramphastos toco (*Ramphastidae* by Federico Degrange, CC0 1.0).

Unique species were themselves diverse, including the Little Spotted Kiwi (*Apteryx owenii*), Pygmy Palm Swift (*Tachornis furcata*), Lesser Frigatebird (*Fregata ariel*), Green Peafowl (*Pavo muticus*), Banded Stilt (*Cladorhynchus leucocephalus*) and Toco Toucan (*Ramphastos toco*). In contrast, 52.6% of birds (5,806 species) had a global uniqueness of <0.1, indicating high overlap with other species, and a high morphological redundancy in the global avian assemblage (230 species have a uniqueness of <0.01). In line with the pervasive pattern of evolutionary convergence in geographically isolated lineages^30^, many species were highly locally unique despite having low uniqueness within the global assemblage (figure 3). At the grid-cell (local) scale there was low morphological redundancy: 24.6% of species (2673 species) had a mean local uniqueness of 1 (averaged across grid cells of species’ current extant range) showing no overlap in trait space with any other birds overlapping their current extant range, and only 230 species had a mean local uniqueness <0.1.

### The functional Red List Index

We calculated the Red List Index (RLI) following the methods outlined in Butchart et al.^16,20^, including 11,044 extant and recently extinct bird species. To calculate the global fRLI we weighted species according to their global uniqueness, and to calculate the local fRLI we weighted species according to their mean local uniqueness across all terrestrial grid cells in their area of habitat. All three indices: RLI, global fRLI and local fRLI showed similar patterns of declines between 1988 and 2024, indicating an increase in aggregate extinction risk over time (figure 4). Across all assessment periods, both the global fRLI and local fRLI were lower than the RLI and showed a slightly greater rate of decline. Uncertainty due to Data Deficient species, species with missing trait data, and species with missing range data, was low for all indices owing to high data availability for birds (figure 4).

**Figure 4.**
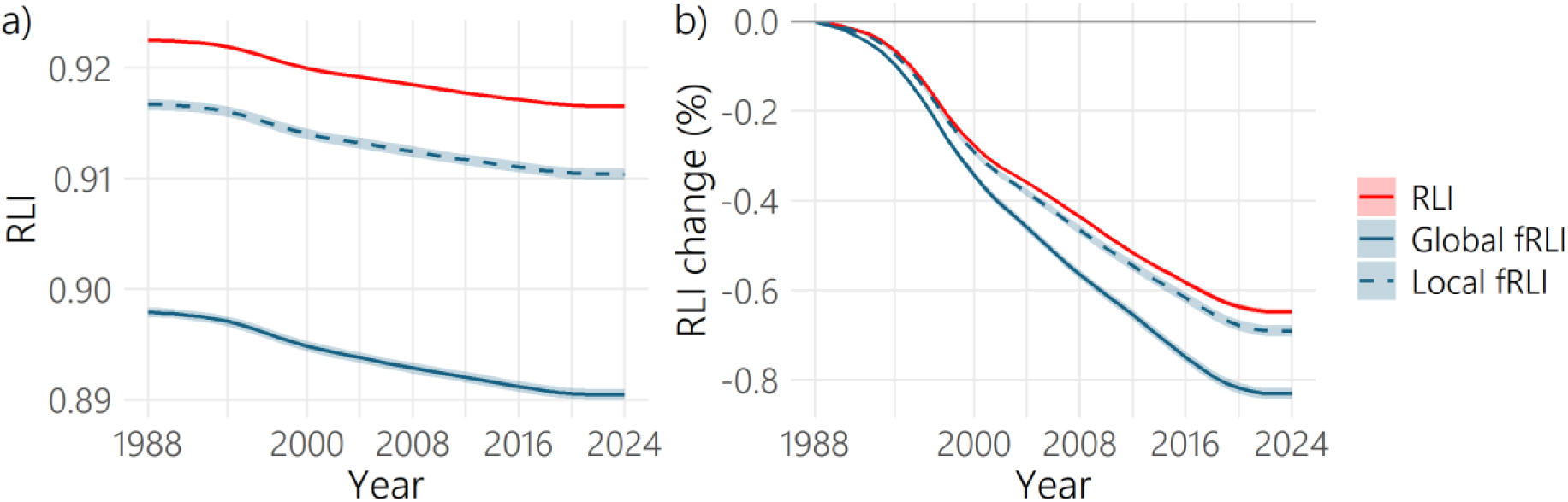
The Red List Index and functional Red List Index between 1988 and 2024. **a,** Red List Index (RLI) and **b,** RLI change between 1988 and 2024, calculated for all birds. Panels show results based on the standard RLI (RLI, all species weighted according to extinction risk), our proposed global functional RLI (Global fRLI, all species weighted according to their extinction risk and global functional uniqueness), and our proposed Local functional RLI (all species weighted according to their extinction risk and mean local functional uniqueness) including 11,044 extant and recently extinct species. Error bands (shading) around the lines represent mean ± 2 x standard deviation of 1,000 simulations where 40 Data Deficient species were assigned extinction risk trajectories at random from species with extinction risk data (of which three species were also missing range data), six species with missing trait data were assigned global functional uniqueness at random from global functional uniqueness of species with trait data, and 179 species with missing range data and six species with missing trait data were assigned local uniqueness at random from local uniqueness of species with trait and range data. A five-year moving average was calculated as described by Butchart et al.^16^ and back-casting was carried out as described in Butchart et al.^20^.

Here we apply the fRLI to birds, but the metric could be applied to any taxonomic group for which there is appropriate coverage of Red List assessments and relevant trait data. As with the RLI^17^, the fRLI could also be disaggregated to show trends for different habitats, ecological groups or regions of interest, and could be adapted to provide national functional Red List Indices^37^. The fRLI provides a way to incorporate functional diversity into conservation planning and policy, to ensure that biodiversity monitoring considers more than only taxonomic diversity^38^.

Limitations that apply to the RLI, such as taxonomic coverage and sensitivity^17^, also apply to the fRLI. The global RLI is currently restricted to mammals, birds, amphibians, corals and cycads, but in the coming decade it will incorporate reptiles, dragonflies, swallowtails, and conifers (based on comprehensive assessments), as well as fishes, freshwater decapods, freshwater molluscs, orthoptera, monocots, legumes, bryophytes, pteridophytes, and fungi (based on assessments of randomized samples^17^). Limitations of taxonomic coverage are compounded when applying the fRLI due to the additional requirement for sufficient trait coverage, for which there is greater availability in better-studied groups such as vascular plants and vertebrates^39,40^. Despite this, the low sensitivity of functional diversity metrics to moderate quantities of missing data^41^ (see Supplementary Information for assessment of sensitivity of uniqueness to missing data) suggests that calculating the fRLI would be practical for many vertebrate groups and some plant groups. Rapidly increasing availability of trait coverage, networks for sharing trait data^42,43^, and the increasing coverage of the RLI^17^ will greatly improve the potential taxonomic coverage of the fRLI in the near future.

### Extinction risk and extinction risk change

To better evaluate factors influencing differences between the RLI and the fRLI, we tested whether globally and locally unique birds had higher or lower extinction risk in 2024 using a phylogenetic generalized linear mixed model (PGLMM) which accounted for non-independence geographically and across the avian tree of life. Species that were more locally unique (β=0.24, pMCMC<0.001, n=10,806 species, figure 5a) or had greater body mass (β= 0.75, pMCMC<0.001, n=10,806 species) had higher extinction risk in 2024 than those that were less locally unique or had lower body mass (figure 5b).

**Figure 5.**
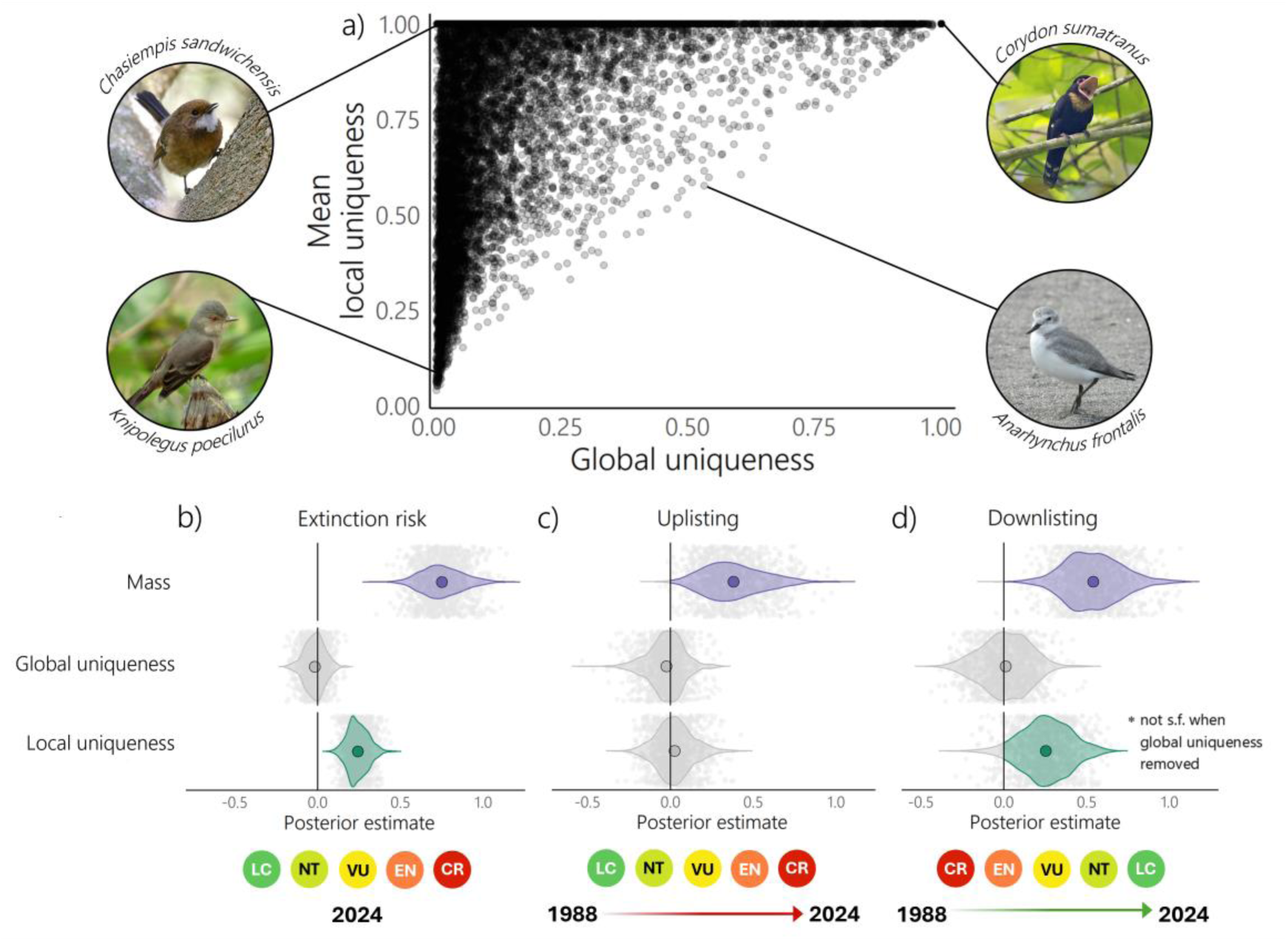
Local uniqueness was positively associated with extinction risk, while global uniqueness was not. **a,** Scatterplot of global uniqueness and mean local uniqueness (n=10,806 species) with examples of species with different combinations of global and mean local uniqueness values. Bottom panels show posterior values from a multi-response Monte-Carlo generalized linear mixed models showing the relationships between body mass, global uniqueness and mean local uniqueness and **b,** extinction risk (n=10,806 species), **c,** the number of categories uplisted between 1988-2024 (uplisting, n=10,756 species), and **d,** the number of categories downlisted between 1988-2024 (downlisting, n=10,494 species). For every model we sampled 1,000 posterior estimates (grey points show individual estimates). Global uniqueness and local uniqueness values were logit transformed, the mean logit-transformed local uniqueness value across species’ area of habitat was calculated, then logit-transformed global uniqueness and mean logit-transformed local uniqueness were centred and scaled to allow comparison of effect size between variables. Mass was log10 transformed and then centred and scaled. Photo attributions: *Chasiempis sandwichensis* by HarmonyonPlanetEarth (<u>CC BY 2.0</u>, photo was cropped and brightened), *Corydon sumatranus* by Lip Kee Yap (<u>CC BY 2.0</u>, photo was cropped), *Knipolegus poecilurus* by Alejandro Bayer Tamayo (<u>CC BY 2.0</u>, photo was cropped), *Anarhynchus frontalis* by Emily Roberts (<u>CC BY 4.0</u>, photo was cropped).

Our finding adds to growing evidence that patterns of species extinction risk are biased with respect to species traits^22,44,45^. However, contrary to previous studies^23,24^, we find that local uniqueness, rather than global uniqueness was important for describing variation in extinction risk. Global uniqueness was not important for explaining variation in extinction risk (pMCMC=0.78, n=10,806 species, removing this predictor did not change the significant effect of other variables). Previous studies analysing the association between global uniqueness and extinction risk did not consider local uniqueness, and when Ali et al.^23^ accounted for phylogenetic non-independence and/or body size, they found no positive association between global uniqueness and extinction risk. No interaction terms were significant in explaining variation in extinction risk (pMCMC>0.1 for all interaction terms under stepwise removal, n=10,806 species).

Loss of unique species is expected to have negative implications for nature’s contribution to people, as unique species make more distinct contributions to ecosystem function^46^ and are more likely to be utilised by people than less unique species (although this study was conducted at a global scale)^25^. Due to decreased functional redundancy at a local scale^38^, increased vulnerability of locally unique species could have more serious and widespread implications for ecosystem function than extinction risk bias towards globally unique species.

We quantified whether globally and locally unique species were more likely to have experienced uplisting (escalation of extinction risk) or downlisting (reduction in extinction risk owing to recovery) between 1988 and 2024. We found that species with greater body mass were more likely to have experienced uplisting in extinction risk category between 1988 and 2024 (β=0.38, pMCMC=0.008, n=10,756 species), while global uniqueness (pMCMC=0.85, n=10,756 species) and local uniqueness (pMCMC=0.85, n=10,756 species) were not significant for explaining variation in uplisting in extinction risk (figure 5c). Species that were more locally unique (β=0.30, pMCMC<0.001, n=10,494 species) and species that had greater body mass (β=0.56, pMCMC<0.001, n=10,494 species) were more likely to have qualified for downlisting to lower extinction risk categories between 1988 and 2024, but we found no effect of global uniqueness (figure 5d). The significance of body mass was not affected if we removed global uniqueness as an explanatory variable, but local uniqueness became non-significant (pMCMC=0.14, n=10,494 species). No interaction terms were significant in explaining variation in uplisting or downlisting of extinction risk category between 1988 and 2024 (pMCMC>0.1 for all interaction terms under stepwise removal, n=10,757 species [uplisting] and 10,494 species [downlisting].

While locally unique species were more likely to be threatened with extinction, we did not find that they were more likely to become closer to extinction over time. This may be promising for the conservation of functional diversity, although questions remain over why locally unique species are more likely to be threatened with extinction if they are not more likely to be experiencing escalations in extinction risk over time. It is possible that locally unique species are more sensitive to change, but not necessarily more prone to decline^47,48^. Alternatively, locally unique species may be more likely to be covered by conservation action, potentially reducing the probability of uplisting and increasing the probability of downlisting. We tested this second hypothesis, analysing protected area coverage specifically, owing to the fact that protected area coverage is the focus of major conservation targets^4^, and there is good data availability on the extent of protected area coverage across the world^49^.

### Protected area coverage

We used data from the World Database on Protected Areas (WDPA)^50^ and the World Database on Other Effective Area-Based Conservation Measures (WDOECM)^51^ to assess whether globally and locally unique species had greater or lesser coverage of their resident distribution within protected areas than less unique species, including resident area of habitat as a covariate. Species with higher mean local uniqueness (β=-0.03, pMCMC=0.04, n=8687 species) or species with larger resident area of habitat (β=-0.21, pMCMC<0.001, n=8687 species) had a lower percentage of their resident range covered by site-based conservation measures (protected areas and OECMs) than those with lower mean local uniqueness or smaller resident area of habitat (figure 6). Species with greater global uniqueness were more likely to have greater protected area coverage when protected area point data were excluded (β=0.03, pMCMC=0.05, n=8687 species), but this was marginally non-significant when point data were included (pMCMC=0.102, n=8687 species). In breeding and non-breeding areas of migratory species’ ranges, species with larger breeding area of habitat had lower protected area coverage than those with smaller breeding area of habitat, but protected area coverage did not vary with local uniqueness or global uniqueness (see Supplementary Information).

**Figure 6.**
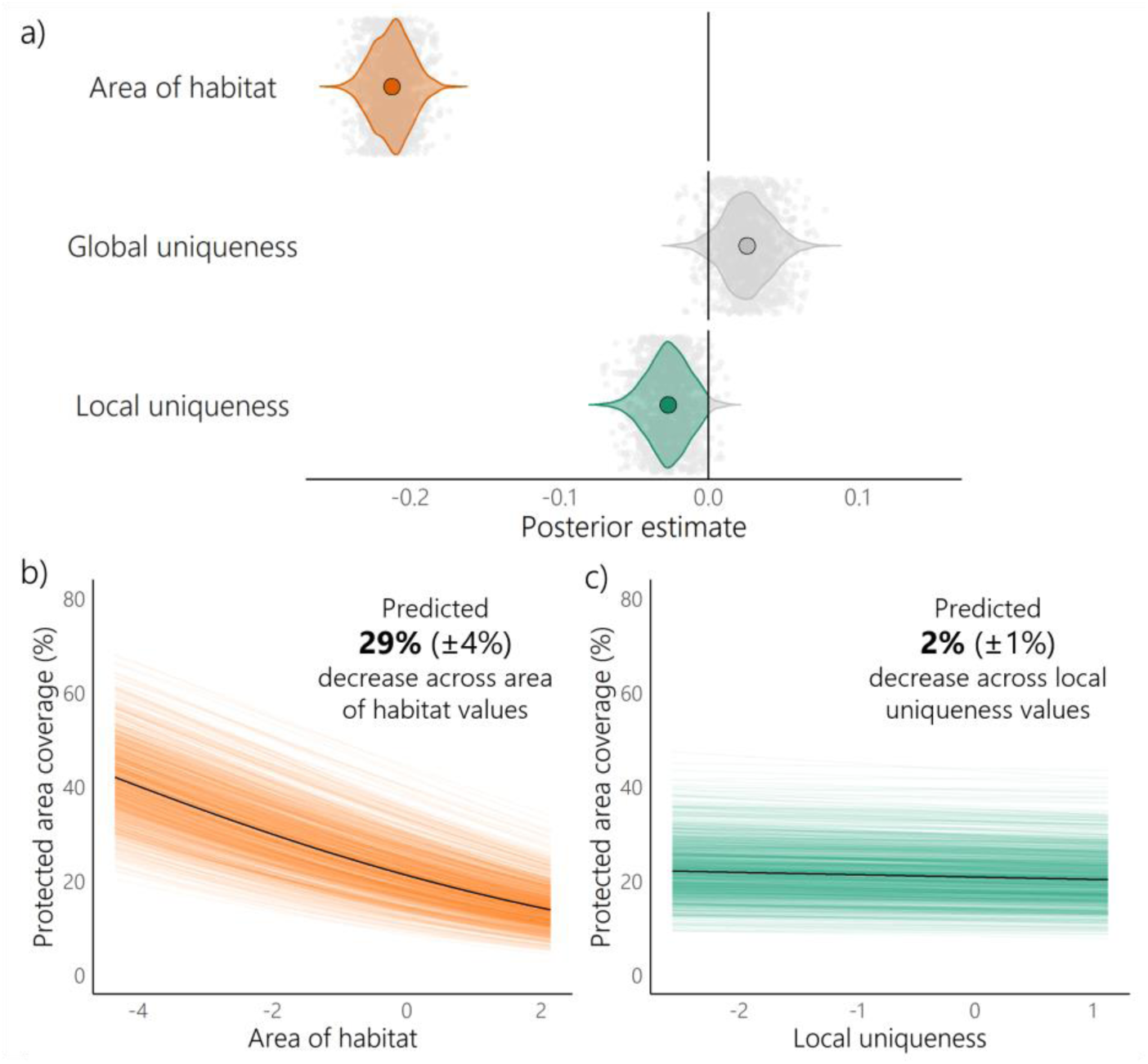
Species with larger area of habitat and greater local uniqueness had lower protected area coverage. **a,** Posterior values from a multi-response Monte-Carlo generalized linear mixed model showing the relationships between area of habitat, global uniqueness and mean local uniqueness and protected area coverage for resident areas of species’ habitats (n=8,687 species). Global uniqueness and local uniqueness values were logit transformed, the mean logit-transformed local uniqueness value across species’ area of habitat was calculated, then logit-transformed global uniqueness and mean logit-transformed local uniqueness were centred and scaled to allow for comparison of effect size between variables. Area of habitat was log10 transformed and then centred and scaled. We sampled 1,000 posterior estimates (grey points show individual estimates). **b,** Predicted protected area coverage across range of scaled log10 transformed area of habitat values of species included in the model (n=8,687 species) for a species with median scaled mean logit-transformed local uniqueness and median scaled global uniqueness (number shows mean difference in protected area coverage with minimum area of habitat and maximum area of habitat ± standard deviation). Orange lines show uncertainty in estimates of model parameters, with different lines representing different model iterations (1,000 posterior estimates). **c,** Predicted protected area coverage across range of scaled mean logit-transformed local uniqueness values of species included in the model (n=8,687 species) for a species with median scaled log10 transformed logit-area of habitat and median scaled global uniqueness (number shows mean difference between protected area coverage with minimum local uniqueness and maximum local uniqueness ± standard deviation). Green lines show uncertainty in estimates of model parameters, with different lines representing different model iterations (1,000 posterior estimates).

Locally unique species were both at greater risk of extinction and less protected from human impacts (although protected area coverage did not vary with uniqueness for breeding and non-breeding areas of migratory species ranges). Existing protected areas have been found to ineffectively safeguard regions that had high risk of losing avian functional diversity in the near future^52^, and poor coverage of functional diversity within protected areas is not restricted to terrestrial ecosystems. In an assessment of the coverage provided by Iberian freshwater protected areas^53^, taxonomic diversity of non-target taxa was well represented but the same protected areas had poor representation of functional diversity. There is evidence that large improvements in the protection of functional diversity could be made with a small increase in protected area coverage^54^. Protecting an additional 5% of land could triple the representation of functional diversity within protected areas, if effectively prioritised^54^. Despite lower coverage within site-based conservation networks, locally unique species were not more likely to experience escalations in extinction risk between 1988 and 2024, and there was some evidence that locally unique species may have experienced greater recovery than less locally unique species during this time (although this was not significant when global uniqueness was not included as an explanatory variable). Our findings highlight the importance of monitoring functional diversity change as the coverage and efficacy of conservation measures can differ between taxonomic and functional diversity.

### Conclusions

The fRLI can be used to monitor the impact of trends in species’ extinction risk on functional diversity, using information on species’ morphological uniqueness and data on genuine changes in extinction risk categories over time. The fRLI for birds shows higher aggregate extinction risk than reflected in the standard RLI. Birds that are unique in their local assemblages (high mean local uniqueness) are more likely to be at high risk of extinction and have poorer coverage of their resident range in protected areas, which could leave them vulnerable to future decline. It is essential to consider functional diversity in every stage of conservation monitoring and practice to provide a comprehensive view of biodiversity change, and the proposed fRLI provides a monitoring tool for this purpose.

## Methods

### Data

We used IUCN Red List data describing current extinction risk categories (as listed in May 2024)^55^ obtained through the R package *rredlist*^56^. IUCN Red List assessments have been carried out for 11,197 bird species (as of May 2024), of which 11,044 species were extant in 1988. Species that went extinct before 1988 were excluded from the analysis.

We obtained data on genuine changes in extinction risk category over the period 1988-2024 from BirdLife International (the Red List Authority for birds, who provide the assessments for all bird species on the IUCN Red List every four to six years). When species are reassessed for the Red List, they may be reclassified in higher or lower categories of extinction risk. Such changes may result from improved knowledge, taxonomic revision, or genuine improvement or deterioration in status (for example, owing to successful conservation actions, or escalating threats). We used data on these genuine changes, excluding reclassifications resulting from changes in knowledge or taxonomy.

We used data on 11 morphological traits from AVONET^27^ including body mass, beak length (two measurements: length from the tip to the skull, and length from the tip to the anterior edge of the nares), beak width, beak depth, wing length, secondary length, Kipp’s distance, hand-wing index, tarsus length and tail length. We included these traits due to their association with avian ecology. Wing morphology describes variation in native dispersal distance (Kipp’s distance^57^, hand-wing index^58,59^), migration behaviour (principal components of wing length and length of primary feathers to the wing tip^60^, Kipp’s distance^57^, isometric wing size^61^, hand-wing index^31^), breeding range size (hand-wing index^31^), and a bird’s ability to fly over unfavoured habitats (hand-wing index^62,63^) and navigate clustered airspace (hand-wing index^31^). Tail size and shape are important for lift, drag and manoeuvrability during flight, affecting a bird’s ability to catch prey in the air, and navigate through forests^64^. Beak morphology is associated with diet and foraging behaviour^30,65,66^, and beak size has implications for thermoregulatory capacity, among other aspects of avian ecology and behaviour^67,68^. Tarsus length is associated with habitat choice^32,69^, and in hot environments, having long legs can assist with thermoregulation^70^.

All 11,044 extant and recently extinct species that were assessed by BirdLife on the IUCN Red List in 2024 were included in the analysis, of which we obtained trait data for 10,990 species, inferred trait data for 48 species (in addition to 860 species which have one or more inferred trait in AVONET), and did not include any trait data for six species (see Supplementary Information). Taxonomic differences between the IUCN Red List and AVONET were resolved using *rl_synonym* in the package *rredlist*^56^ (292 species) and by using trait data captured in AVONET under eBird and BirdTree nomenclature rather than the BirdLife taxonomy used by AVONET version 7 (16 species).

### Functional uniqueness

We described functional uniqueness of species using trait probability densities following the method of Stewart et al.^34^ (figure 2, equation 1). We first summarised variation in the 11 morphological traits (log10 transformed, centred, and scaled to unit variance) using principal component analysis^41^. We selected the first four principal components to give detailed information about species’ ecological niches^30^. Morphological variables were used as they can provide more in-depth information about an organism’s niche than categorical ecological variables^30^.

Species’ probability distributions describe the relative abundance of trait values (in this case principal component values) within a species. Species’ probability distributions were gaussian distributions constructed using the R package *TPD*^33^ where the means were principal component values derived from functional trait data^27^ and standard deviation was estimated using a bandwidth selector (*Hpi.diag* function from package *ks*^36^). Sensitivity analyses revealed that our results were robust to deviation in estimated standard deviations of traits (Supplementary figure 1 and 2) but nevertheless do not account for variation in intraspecific trait variance across species. Community probability distributions were constructed by calculating the sum of species’ probability distributions and describe the relative abundance of trait values across the community (global or local assemblage). Trait space was divided into 390,625 cells (25 bins for each principal component) to facilitate construction of species’ probability distributions and community probability distributions^33^.

The uniqueness metric (equation 1) describes the probability that functional richness is reduced when a species goes extinct. Uniqueness was computed for each species by calculating the density of the species’ probability distribution relative to the community probability distribution for all cells where the species’ probability distribution was greater than 0. Areas of greater relative species’ probability distribution density were weighted to give greater contribution to species’ uniqueness values. This uniqueness metric ranges between 0 and 1, where values close to 0 indicate that the species occupies the same area of trait space as many other species and so their species probability distribution overlaps with the species probability distribution of many other species in gridded trait space. A value of uniqueness close to 0 indicates that extinction is unlikely to result in functional richness loss. On the contrary a value of 1 indicates that the species occupies an area of trait space that is not shared with any other species, with no overlap between the species probability distribution and the species probability distributions of other species, indicating that extinction will lead to functional richness loss.

### Global and local uniqueness

Species may be unique at a global scale (very different morphologically from other extant species) or unique at a local scale (very different morphologically from other species across their range and habitat, even if potentially similar to other species living elsewhere). To capture both scales we defined *global uniqueness* as the functional uniqueness of species within the global assemblage, and *local uniqueness* as the functional uniqueness within local terrestrial assemblages. When calculating global uniqueness, the community probability distribution was built with all 11,038 species for which we had measured or inferred trait data. While only six bird species are missing trait data, data availability can be far more limited in other taxa and could impact the applicability of our approach to other groups. We tested the impact of missing data on global uniqueness estimations, comparing the metric proposed here as well as another recently proposed metric for identifying functionally unique species, as part of the functionally unique, specialised and endangered species (FUSE) metric (as applied by Pimiento et al.^72^) (Supplementary figure 3). Both the FUSE uniqueness metric and functional uniqueness calculated here were resilient to 30% missing data with random and biased removal of data. However, our functional uniqueness metric had less error in functional uniqueness estimation in most missing data scenarios. Both metrics had high error when 70% of data were missing, so we recommend imputing missing data where possible^41^ and using caution when applying the fRLI to highly incomplete datasets.

To calculate local uniqueness, we generated an equal area square grid with 110 km sides (grid cell area: 12,100 km^2^) in the Behrmann projection^73^. We used area of habitat maps from BirdLife International^35^ (building on the approach of Lumbierres et al.^74^) in the Behrmann projection^73^ at a resolution of 1km^2^, which allowed us to exclude areas of species’ ranges where species were highly unlikely occur (proportional suitability of less than 0.01) due to known habitat and elevation preferences. We identified grid cells occupied by species across their resident, breeding and non-breeding area of habitat.

Where area of habitat maps showed no suitable habitat was present (22 species), or for species where the taxon identification number in the IUCN Red List^55^ was not found in the area of habitat maps (425 species, either due to taxonomic updates or insufficient data availability), we used species’ range boundaries from BirdLife International and Handbook of the Birds of the World^75^. For seven species, only breeding or non-breeding distributions were missing from the area of habitat maps, so we used range polygons for the appropriate season, while for the rest (440 species) we selected parts of the species’ range where the seasonality was coded as resident, breeding, non-breeding or seasonality uncertain (i.e. excluding passage). We then selected range polygons where the origin was coded as native, reintroduced, or assisted colonisation (i.e. excluding introduced, vagrant and origin uncertain), and where the presence was coded as extant (i.e. excluding possibly extant, possibly extinct, extinct, presence uncertain, and expected additional range where the species is now extinct), in line with the approach used for area of habitat maps^74^. We merged selected species’ range polygons using the function *st_union* in the *sf* package^76^. We transformed the Copernicus Global Land Service Dynamic Land Cover layer for 2019 at 100m resolution^77^ to Behrmann projection^73^, selected areas of the raster to exclude sea pixels (VALUE<200), converted the raster to polygons, and merged polygons to provide a land/sea mask. We clipped the merged species’ range polygons by the land/sea mask, then conducted a spatial join with the grid to give a list of species whose range overlapped with each grid cell.

We combined the list of species that occupied each grid cell calculated using area of habitat maps with that calculated using range maps, to define “local communities”. We excluded grid cells that only included species whose distribution data was obtained from range polygons (terrestrial communities within predominantly marine cells), as this was an artefact of the high resolution of the land/sea mask (100m resolution) relative to resolution of the area of habitat maps (1km resolution). For each grid cell, we built a community probability distribution, and calculated species uniqueness for all species present in the grid cell. We finally calculated mean local uniqueness for all species, across all grid cells where they were present. For 179 species it was not possible to calculate local uniqueness because they had no terrestrial range (145 species, including those restricted to small islands or ice shelves and sea ice), or no known remaining extant range (34 species). Spatial analyses were conducted in ArcGIS Pro^78^ and R^79^.

### Species and Functional Red List Index

We calculated the Red List Index (RLI) according to Butchart et al.^20^ (equation 2), with 11,044 extant and recently extinct species.

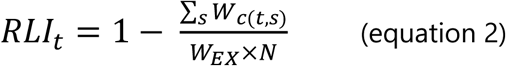

Where, *RLI*_*t*_ is the index value in assessment period *t*, *W*_*c*(*t*,*s*)_ is the weight of the IUCN Red List Category of species *s* in assessment period *t*, *W*_*EX*_ is the weight given to extinct species, and *N* is the number of species that were not extinct in 1988, the start of the earliest assessment period. We used equal-steps weights as recommended by Butchart et al.^20^, meaning Least Concern was given a weight of 0, Near Threatened a weight of 1, Vulnerable a weight of 2, Endangered a weight of 3, Critically Endangered a weight of 4 and Extinct — including species listed as Critically Endangered (Possibly Extinct), Extinct in the Wild, and Critically Endangered (Extinct in the Wild) — a weight of 5.

Following the RLI method, we assumed that the current Red List category applied in every assessment since the first, except for those species documented as having undergone genuine changes in category (i.e. we back-cast Red List categories using those published in May 2024^55^, except for those that had undergone genuine changes). This means that non-genuine changes resulting from improved knowledge or taxonomic changes do not impact estimations of the RLI^20^.

A uniqueness weighting was obtained by dividing the uniqueness of a given species, by the summed uniqueness values for all species, and multiplying by the number of species (equation 3). Uniqueness weightings were calculated separately based on the species’ global uniqueness and mean local uniqueness across a species’ range.

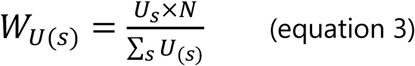

Where *W*_*U*(*s*)_ is the uniqueness weighting of species *s*, *U*_*s*_ is the uniqueness value of species s and *N* is the number of species.

The fRLI was calculated in the same way as the RLI, but each species was multiplied by its uniqueness weighting (equation 4). In the global fRLI the global uniqueness weighting for each species was used, and in the local fRLI the local uniqueness weighting was used.

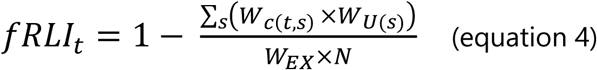

Where, *fRLI*_*t*_ describes the fRLI in assessment period *t*, *W*_*c*(*t*,*s*)_ is the weight of the IUCN Red List Category of species *s*, in assessment period *t*, *W*_*U*(*s*)_ is the uniqueness weighting of species *s*, *W*_*EX*_ is the weight given to extinct species, and *N* is the number of species that were not extinct in 1988, the earliest assessment year.

In both the RLI and fRLI there is temporal uncertainty because comprehensive reassessments are conducted over multi-year periods (every six years before 2000 and every four years after 2000). If a species showed a genuine change during an assessment period, this change could have occurred in any year of that assessment period. To reflect this, RLI and fRLI values were interpolated linearly between assessment periods, and a moving average with a window of five years (three years for the first and last year, and four years for the second and penultimate year) was calculated, following Butchart et al.^16^.

To quantify the impact of uncertainty due to missing extinction risk data (40 Data Deficient species), missing trait data (six species, of which four are now extinct) and missing range data (179 species), we generated 1,000 datasets from which we calculated the RLI, the global fRLI and the local fRLI. For each dataset we assigned Data Deficient species with a trajectory of extinction risk categories selected at random from data sufficient species (a trajectory is the entire sequence of risk categories over the study period). This approach assigns extinction risk categories to Data Deficient species that reflect the proportion of categories in the data within assessment periods, while ensuring temporal consistency between assessment periods (reflecting observed trajectories). Ensuring we assigned observed trajectories was important because species are weighted unequally to reflect differences in uniqueness and the index could be affected if a unique species had an extreme trajectory which is unobserved in the dataset, such as being downlisted from CR to LC over one assessment period. Each species with missing trait data was assigned a global uniqueness value selected at random from the global uniqueness values of species with complete trait data. Species with missing range data and species with missing trait data were assigned a mean local uniqueness at random from the mean local uniqueness values of species with complete trait and range data. This was repeated 1,000 times, and the mean and standard deviation of the RLI and fRLI values were calculated.

### Modelling extinction risk

We modelled extinction risk against global and local uniqueness to find whether more unique species had a greater or lesser extinction risk than less unique species. Data Deficient species (40 species), species with missing trait data (6 species) and species for which it was not possible to calculate local uniqueness (179 species, of which three were Data Deficient) were removed. A further 16 species that have been recently described and have no suitable match in Jetz et al.^80^ were also removed. The final models were fitted for 10,806 species.

We used a Markov chain Monte Carlo multivariate generalised linear mixed model (MCMCglmm), with extinction risk as the ordinal dependent variable and global uniqueness (continuous, logit transformed then centred and scaled), mean local uniqueness (continuous, logit transformed then centred and scaled), and body mass (continuous, log10 transformed then centred and scaled), as independent variables. Body mass was included as a known correlate of extinction risk. Local uniqueness for each species in each occupied grid cell was logit transformed, then the mean of logit-transformed local uniqueness values for each species was calculated. All explanatory variables were centred and scaled (normalised) to allow for comparison of effect sizes between variables. Initially all interactions between body mass, global uniqueness and mean local uniqueness were included. If found to be non-significant, the three-way interaction term was removed. Then, we refitted the model, and removed the least significant two-way interaction variable, if any were non-significant. We repeated until only significant interactions remained.

Phylogeny and spatial variables were included as random effects to account for non-independence between species due to shared evolutionary history and spatial autocorrelation. For the phylogenetic random effect, we used a maximum clade credibility tree constructed from the first 1,000 trees under the Hackett backbone of Jetz et al. (2012). Where species had two synonyms under BirdTree, the synonym with the exact species name match was chosen (98 species). A further four BirdLife and BirdTree synonym pairs were matched that had the same species name but different suffixes to reflect the gender of the genus name (for example, *Psittacara holochlorus* [BirdLife] was matched to *Aratinga holochlora* [BirdTree]). For two species, synonyms in BirdTree were picked at random. For 48 species which were not present in AVONET under any taxonomy, the tip of the closest extant relative was used, identified using the Handbook of the Birds of the World and BirdLife International digital checklist of the birds of the world^81^ (in the same way that the closest extant relatives were identified for inferring trait data). Where two or more species under the BirdLife taxonomy had one synonym in BirdTree, the same tip from the maximum clade credibility tree was used to represent more than one species (842 tips represented more than one species under the BirdLife taxonomy).

We accounted for spatial non-independence using eigenvectors of the centroids of species breeding and resident ranges including areas where the species was coded as extant and either native or reintroduced. Centroids were obtained from AVONET^27^ where available, and calculated following the method of Tobias et al.^27^ for 81 species (including 69 species for which trait data was inferred or obtained from AVONET under the eBird or BirdTree taxonomies, and 12 species which were missing data on range centroid longitude and latitude). Eigenvectors of centroids were calculated using principal coordinates analysis, implemented through the *ape* package^82^ on a Haversine distance matrix between species centroids, calculated using the *sf* package^76^. Four spatial eigenvectors provided optimal description of variance in distances between species centroids (identified through scree plots, Supplementary figure 6) so were included as random effects.

MCMCglmms were fitted using the R package *MCMCglmm*^83^. MCMC chains were run for 200,000 iterations with a 100,000-iteration burn-in period and a thinning interval of 100 iterations for models. Model convergence was assessed using plots of parameter traces produced through the *plot* function in package *MCMCglmm*^83^. We used a chi-squared prior for the phylogenetic random effect as this best approximates a uniform distribution^84^. Cauchy-scaled Gelman priors were used for fixed effects, as is recommended by Gelman^85^ for ordinal regressions. It is not possible to estimate residual variance with an ordinal dependent variable, so the residual variance was fixed at 1 ^86^. For the phylogenetic random effect, we used a chi-squared prior (expected covariance of 1, degree of belief of 1000, mean vector of 1, and covariance matrix of 1), as this best approximates a uniform distribution, giving an uninformative prior^84,86,87^. For spatial random effects we used parameter expanded priors (expected covariance of 1, degree of belief of 1, mean vector of 0, and covariance matrix of 625) as they are often less informative than the default inverse-Wishart prior^86^.

Significance was assessed using the pseudo p-value (pMCMC) provided by the *MCMCglmm* function in package *MCMCglmm*^83^. A significance threshold of 0.1 rather than 0.05 was used, to give a one-tailed significance test that the posterior distribution overlaps with zero, rather than the default two-tailed significance test^88^.

### Modelling extinction risk change

To assess whether globally and locally unique species were more likely to experience uplisting or downlisting in extinction risk we fitted two models: one considering only species that showed no change or were uplisted between 1988 and 2024 (n=10,764 species), and another considering only species that showed no change or were downlisted (n=10,494 species), with global uniqueness (continuous, logit transformed then centred and scaled), mean local uniqueness (continuous, logit transformed then centred and scaled), and body mass (continuous, log10 transformed then centred and scaled) as explanatory variables. All explanatory variables were centred and scaled to allow for comparison of effect sizes between variables. Change in extinction risk was an ordinal factor describing the number of extinction risk categories that a species was uplisted or downlisted by between 1988 and 2024. The minimum possible value of change in extinction risk was no change, while the maximum possible value was an uplisting or downlisting of four categories (from Least Concern to Critically Endangered or Critically Endangered to Least Concern). Initially all interactions between body mass, global uniqueness and mean local uniqueness were included. If found to be non-significant, the three-way interaction term was removed. Then, we refitted the model, and removed the least significant two-way interaction variable, if any were non-significant. We repeated until only significant interactions remained.

As above, phylogeny was included as a random effect with a chi-squared prior^84^, and four spatial eigenvectors were included as random effects with parameter expanded priors^86^. We also included the extinction risk category in 1988 as a numerical random effect (using equal weights where 1 indicated Least Concern and 4 indicated Critically Endangered) with a parameter expanded prior, to account for the fact that species may be more or less likely to be uplisted or downlisted based on their extinction risk category at the start of the time period. Cauchy-scaled Gelman priors were used for fixed effects. MCMC chains were run for 200,000 iterations with a 100,000-iteration burn-in period and a thinning interval of 100 iterations.

### Modelling protected area coverage

We aimed to assess whether globally and locally unique species had greater or lesser coverage of their distributions by protected areas than less unique species. To do this we calculated the percentage of each species’ area of habitat (or range where area of habitat data were not available) that was covered by protected areas or Other-Effective Area-Based Conservation Measures (OECMs), using data from the World Database on Protected Areas (WDPA)^50^ and the World Database on Other Effective Area-Based Conservation Measures (WDOECM)^51^. We used the *wdpar* package^89^ to clean WDPA and WDOECM data^90–92^. We used publicly available data on protected areas and Other Effective Area-Based Conservation Methods, including sites which had a status of "*Designated*", "*Inscribed*", "*Established*" or "*Adopted*", and excluding “*Proposed*” sites or sites where the status was not reported. As recommended by Protected Planet^91^, UNESCO Man and Biosphere Reserves (MAB) were removed, as MAB sites include buffer and transition zones that are often not protected areas. Point data without a reported area were also excluded. Remaining sites in the WDPA and WDOECM were reprojected to the Behrmann projection^73^.

Of area-based conservation sites within the World Database on Protected Areas (WDPA)^50^ and the World Database on Other Effective Area-Based Conservation Measures (WDOECM)^51^ 4% have a known area of extent and point location, but do not have a reported polygon defining the boundaries of the protected area. For these sites we fitted a circular buffer that was the size of the reported area around the point location. As buffered centroids are expected to only roughly overlap with actual protected area boundaries^93^ we conducted the analyses both with and without protected area point data (Supplementary figure 4).

Protected area geometries were simplified with a tolerance of 10m to reduce computational burden, and slivers with an area of less than 0.1 square meters were removed. We used a high geometry precision of 10,000 ^89^. Finally, we corrected errors in the geometries using the *st_make_valid* function in package *sf* ^76^, and dissolved geometries into a single geometry, referred to as the protected area layer.

We calculated the total habitat area where species were resident by selecting cells in resident area of habitat maps with a proportional suitability greater than 0.01. We then calculated the area of resident habitat (proportional suitability > 0.01) covered by the protected area layer by calculating the sum of coverage area across all cells in each area of habitat map using the R package *exactextractr*^94^. Protected area coverage was expressed as a percentage of the total resident habitat area. We repeated this for non-breeding and breeding areas of habitat (migratory species only) separately, as protection from threats across all stages of the annual cycle is important for the conservation of migratory species^94^ (see Supplementary Information).

For species that had no suitable habitat present (22 species), or for species where the taxon identification number in the IUCN Red List^55^ was not found in the area of habitat maps (425 species, either due to taxonomic updates or insufficient data availability), we calculated the merged polygons of each species’ range as described in the *Global and local uniqueness* section of the methods, but separated resident (including seasonality uncertain), breeding and non-breeding areas of species’ ranges. For each species and season, we calculated the area of the intersection between the protected area layer and the merged species range map, and reported protected area coverage as a percentage of the total area of the range polygon for a given species and season.

We fitted gaussian MCMCglmm models with protected area coverage (continuous, logit transformed) as the response variable and global uniqueness (continuous, logit transformed the centred and scaled), mean local uniqueness (continuous, logit transformed then centred and scaled) and area of habitat (log10 transformed then centred and scaled) as independent variables, for resident (n=8687 species), breeding (migratory species only, n=2297 species) and non-breeding (migratory species only, n=2201 species) area of habitats (and species ranges) separately. Area of habitat was included to control for its expected relationship with protected area coverage. Phylogeny was included as a random effect with a chi-squared prior^84^. When modelling protected area coverage, the fourth spatial eigenvector had low posterior variance (indicating it was not useful for explaining covariance between species) which resulted in singularities when fitting the model, so it was removed. Three spatial eigenvectors were included as random effects with parameter expanded priors^86^. Normal priors were used for fixed effects, with an expected value (μ) of 0 and strength of belief (V) of 1 x 10^8 86^. Due to singularities in the model of breeding and non-breeding protected area coverage for some random seeds, we compared models run with and without spatial random effects, which did not change our conclusions (see Supplementary Information). MCMC chains were run for 103,000 iterations with a 3000-iteration burn-in period and a thinning interval of 100 iterations, as they required less time to converge than ordinal models.

## Data availability

AVONET data on morphological, ecological and geographical traits for all birds**^Error! Reference source not found.^** is available for use under a creative commons licence (<u>CC BY 4.0</u>): https://doi.org/10.6084/m9.figshare.16586228.v7. Data on IUCN extinction risk categories, and threats affecting each species, are available from the IUCN Red List^55^ and can be accessed through the package rredlist^56^. Information on terms of use of IUCN Red List data can be found at https://www.iucnredlist.org/terms/terms-of-use. Data on genuine changes in extinction risk, species’ area of habitat maps and species’ distribution maps can be requested from BirdLife (https://datazone.birdlife.org/dataset-information). The World Database on Protected Areas (WDPA)^50^ and the World Database on Other Effective Area-Based Conservation Measures (WDOECM)^51^ are available for non-commercial use and more information on terms of use can be found at https://www.protectedplanet.net/en/legal.

## Code availability

Code will be made publicly available under a creative commons license upon acceptance.

