## Supplementary Information for "A functional Red List Index for monitoring trends in functional diversity"

#### Obtaining trait data for all species with extinction risk assessments

Trait data were obtained from AVONET (Tobias et al., 2022) and inferred from closely related species where trait data were not available. AVONET provides trait data under three taxonomies: the BirdLife taxonomy (Handbook of the Birds of the World and BirdLife International, 2024), the eBird taxonomy (Clements et al., 2021), and the nomenclature used by Jetz et al. (2012), referred to as the BirdTree taxonomy. Of 11,044 extant and recently extinct species assessed by BirdLife on the IUCN Red List in 2024, 10,686 species had a direct match to the 2020 BirdLife taxonomy in AVONET. Of the remaining 358 species, 292 taxonomic differences (resulting from the adoption of taxonomic revisions by BirdLife in the intervening years) were resolved using the function *rl_synonym* in the package *rredlist* (Chamberlain, 2020). A total of 16 species synonyms were captured in AVONET under different taxonomies (11 synonyms under eBird, and 5 under BirdTree). For these synonyms, and the species with which they were synonymous, trait data were obtained from AVONET under the BirdTree or eBird taxonomy. Of 10,990 species for which trait data were obtained from AVONET, 860 species (7.8%) had inferred trait data for at least one trait. In addition to these species, we inferred trait data for another 48 species from the closest extant relative present in AVONET, identified using the Handbook of the Birds of the World and BirdLife International digital checklist of the birds of the world (Handbook of the Birds of the World and BirdLife International, 2024). For two species, *Heliothraupis oneilli* and *Gallirallus lafresnayanus*, there was not an appropriate close relative to infer trait data from. *Heliothraupis oneilli* was first described in 2021 and has been classified in its own genus (Lane et al., 2021), and *Gallirallus lafresnayanus* has no trait data because there have only been two unconfirmed sightings in the last 100 years (BirdLife International, 2024a), and its taxonomic relationships and closest relatives are uncertain. As a result, 11,044 extant and recently extinct species were included in the analysis, of which we obtained trait data for 10,990 species, inferred trait data for 48 species, and did not include any trait data for six species.

#### Intraspecific variation

To construct species probability densities, data is required on the mean of traits for each species and on intraspecific variation in species traits, namely the standard deviation in trait values. Where good data for intraspecific trait variability is lacking, the standard deviation of traits can be estimated using a bandwidth selector (see Methods), as done by Carmona et al. (2021), Llorente-Culebras et al. (2024) and Toussaint et al. (2024).

AVONET contains unparallelled data on intraspecific trait variability, with complete measures for at least four individuals for almost 9000 species (8072 under the BirdLife taxonomy). Nevertheless, data availability for other taxonomic groups is far lower than for birds, and we intended the functional Red List Index to be applicable to other taxonomic groups. As such, when calculating uniqueness, we estimated intraspecific trait variability using a bandwidth selector (*Hpi.diag* function from package *ks* [Duong, 2021]).
We tested the impact of deviation from estimated standard deviation in traits. We increased and decreased the standard deviation of traits by 20% when constructing species probability distributions and calculated uniqueness.

There was high agreement between estimates of global uniqueness when calculated with different degrees of intraspecific trait variation (Supplementary figure 1). Global uniqueness with high trait variability (+20% of that estimated by the bandwidth selector) explained 89% of the variance in global uniqueness values (linear model of global uniqueness against global uniqueness with increased trait variability, $\beta=1.01$, $R^2 = 0.89$, $p<0.0001$, n=11 038 species), while global uniqueness with low trait variability (-20% of that estimated by the bandwidth selector) explained 85% of the variance in global uniqueness values (linear model of global uniqueness against global uniqueness with decreased trait variability, $\beta=0.86$, $R^2 = 0.85$, $p<0.0001$, n=11 038 species).

Likewise, there was high agreement between estimates of local uniqueness when calculated with different degrees of intraspecific trait variation (Supplementary figure 1). Local uniqueness with high trait variability (+20% of that estimated by the bandwidth selector) explained 86% of the variance in local uniqueness values (linear model of local uniqueness against local uniqueness with increased trait variability, $\beta=0.94$, $R^2 = 0.86$, $p<0.0001$, n=10 859 species), while global uniqueness with low trait variability (-20% of that estimated by the bandwidth selector) explained 82% of the variance in local uniqueness values (linear model of local uniqueness against local uniqueness with decreased trait variability, $\beta=0.91$, $R^2 = 0.82$, $p<0.0001$, n=10 859 species).

Estimates of the functional Red List Index (fRLI) were similar whether calculated with estimated, increased or decreased trait variability (Supplementary figure 2).

#### Functional uniqueness methods comparison

Before calculating the functional Red List Index (fRLI), we compared two metrics which can be used to quantify functional uniqueness within the global assemblage (global uniqueness); the Functionally Unique, Specialised, and Endangered metric (FUSE) (Pimiento et al., 2020), and our own metric based on trait probability densities (equation 1 in the main text, referred to here as PRUNE to describe probabilistic uniqueness). The FUSE metric is faster to compute than PRUNE, but PRUNE has the advantage that it is readily interpretable, quantifying the probability of functional richness loss if a given species is removed from a community. For birds we have near complete trait data, however if the fRLI is to be applied to other taxonomic groups it is likely that trait data will be incomplete. Therefore, we compared the response of FUSE and PRUNE to incomplete datasets. This is particularly important as both uniqueness metrics rely on quantifying species positions in trait space relative to other species in the community, so if some species are missing trait data it will also affect the uniqueness values of species for which there is complete trait data.

We tested levels of incompleteness in trait datasets, removing all trait data for 30% or 70% of species. We tested biased and random removal of data. In the random scenario species trait data were removed at random. In the biased scenario, the probability that species trait data were removed from species with a low body mass was elevated, such that the probability that data were removed from species with a body mass less than the median body mass was 0.8 and the probability of removal for species with a body mass over the median body mass was 0.2. This reflects biases of data completeness observed in empirical trait datasets (González-Suárez et al., 2012; Stewart et al., 2023). 100 iterations were run for each of the four scenarios (random removal of data from 30% of species, random removal of data from 70% of species, biased removal of data from 30% of species, and biased removal of data from 70% of species).

For each of the 400 datasets with data removal we calculated FUSE and PRUNE for every remaining species in the dataset. For PRUNE, functional uniqueness was calculated as described in the main text. For FUSE, functional uniqueness was calculated according to Pimiento et al. (2020), by calculating the mean Euclidean distance to the five nearest neighbours of a species. We compared uniqueness values to those estimated with the complete dataset (no data removal), and calculated the mean normalised root mean square error of uniqueness values for every incomplete dataset (normalised by the standard deviation of uniqueness values for each metric).

We used a three-way ANOVA to find how error in functional uniqueness estimations compared between metrics, for scenarios with different proportion of missing data, and random or biased removal. We included two- and three-way interactions between variables (metric, completeness and bias). All terms were significant (n=800, $p<0.001$ for all terms). The percentage of species with missing trait data had the largest effect size (F=9579.72) followed by the metric used (F=499.35). We conducted a Tukey-HSD to find which metric under which completeness and bias scenarios had lowest error in uniqueness estimation.

In nearly all scenarios, PRUNE had lower error in functional uniqueness estimates when using incomplete datasets (Supplementary figure 3, Supplementary table 2). The only exception was that FUSE had lower NRMSE when 70% of species were missing data, and data removal was biased. However, under this scenario both FUSE and PRUNE had high error, so we recommend that if there is a high proportion of missing trait data, the fRLI should not be calculated unless accurate imputation is used (Stewart et al., 2023).

### Protected area coverage in resident area of habitat with and without WDPA points

Of reported area-based conservation sites within the World Database on Protected Areas (WDPA) (UNEP-WCMC, 2024a) and the World Database on Other Effective Area-Based Conservation Measures (WDOECM) (UNEP-WCMC and IUCN, 2024), 4% are reported as point data, either because the data providers lack data on the protected area boundaries or because the information is politically sensitive (Bingham et al., 2019). For points with reported area, we fitted a circular buffer around protected area centroids to match their spatial extent, however as buffered centroids are expected to only roughly overlap with actual protected area boundaries this can introduce error into estimates of species protected area coverage (Visconti et al., 2013). We therefore repeated our analyses to assess whether removing point data, rather than including buffered centroids, affected our conclusions.

Species with larger resident area of habitat (with points: $\beta=-0.21$, $pMCMC<0.001$, n=8687 species; without points: $\beta=-0.22$, $pMCMC<0.001$, n=8687 species), or higher mean local uniqueness (with points: $\beta=-0.03$, $pMCMC=0.04$, n=8687 species; without points: β=-

0.04, pMCMC<0.001, n=8687 species) had a lower percentage of their resident range covered by site-based conservation measures (protected areas and OECMs) than those with smaller resident area of habitat or lower mean local uniqueness whether points were included or excluded (figure 5, Supplementary figure 4). Species which had greater global uniqueness were more likely to have greater protected area coverage when point data were excluded (β=0.03, pMCMC=0.05, n=8687 species), but this was marginally not significant when point data were included (pMCMC=0.102, n=8687 species) (Supplementary figure 4).

#### Protected area coverage in breeding and non-breeding area of habitat

We modelled protected area coverage for non-breeding and breeding areas of habitat separately, as protection from threats across all stages of the annual cycle is important for the conservation of migratory species (UNEP-WCMC, 2024b). Migratory species do not necessarily have both breeding and non-breeding areas of habitat, which is particularly the case for species which are partially migratory, and for some migratory species breeding or non-breeding areas of habitat are not known.

Gaussian MCMCglmm models were run as described in the main text but when modelling protected area coverage of non-breeding ranges, we calculated range centroids from unions of the parts of species ranges that were non-breeding, resident or had uncertain seasonality (areas of species ranges that are included in non-breeding area of habitat maps, BirdLife International, 2024b). When modelling breeding and non-breeding protected area coverage, we calculated eigenvectors of centroids of species ranges included in each model (2,297 and 2,201 species respectively). Due to singularities in the model of breeding and non-breeding protected area coverage for some random seeds, we compared models run with and without spatial random effects. We also compared results when including buffered centroids of point data with reported area in the WDPA and WDOECM, and when removing point data.

In breeding areas of species' ranges, species with larger breeding area of habitat (β=-0.48, pMCMC<0.001, n=2,297 species for all models, Supplementary figure 5) had lower protected area coverage than those with smaller breeding area of habitat, but protected area coverage did not vary with local uniqueness (pMCMC=0.69-0.72, n=2,297 species) or global uniqueness (pMCMC=0.94-0.97, n=2,297 species). In non-breeding areas of migratory species' ranges, species with larger breeding area of habitat (β=-0.25, pMCMC<0.001, n=2,201 species for all models) also had lower protected area coverage than those with smaller breeding area of habitat, and protected area coverage did not vary with local uniqueness (pMCMC=0.21-0.25, n=2,201 species) or global uniqueness (pMCMC=0.12-0.21, n=2,201 species) (Supplementary figure 5). Including or excluding spatial random effects, or including or excluding WDPA and WDOECM, did not affect our conclusions (Supplementary figure 5).

### Supplementary Tables

**Supplementary Table 1** Trait loadings of principal components. The first four principal components were used as they have been shown to accurately predict a bird's ecological niche (Pigot et al., 2020).

| | PC1 | PC2 | PC3 | PC4 | PC5 | PC6 | PC7 | PC8 | PC9 | PC10 | PC11 |
|---|---|---|---|---|---|---|---|---|---|---|---|
| **Cumulative Variance Explained (%)** | 0.666 | 0.816 | 0.893 | 0.937 | 0.972 | 0.981 | 0.99 | 0.996 | 0.998 | 1 | 1 |
| **Mass** | 0.347 | 0.056 | -0.106 | 0.118 | -0.316 | 0.399 | -0.642 | -0.046 | -0.374 | 0.185 | 0.067 |
| **Wing Length** | 0.348 | -0.140 | -0.242 | 0.083 | 0.029 | 0.345 | 0.236 | -0.035 | 0.077 | -0.676 | 0.394 |
| **Tarsus Length** | 0.284 | 0.322 | -0.273 | 0.389 | -0.396 | -0.638 | 0.107 | 0.075 | -0.058 | -0.074 | 0.028 |
| **Beak Length Nares** | 0.295 | 0.003 | 0.601 | 0.222 | 0.223 | -0.037 | 0.254 | -0.089 | -0.574 | -0.125 | -0.184 |
| **Beak Length Culmen** | 0.315 | 0.043 | 0.464 | 0.381 | 0.113 | 0.039 | -0.173 | 0.113 | 0.641 | 0.179 | 0.195 |
| **Beak Width** | 0.319 | 0.109 | 0.176 | -0.590 | -0.135 | -0.060 | 0.027 | 0.694 | -0.023 | -0.045 | 0.026 |
| **Beak Depth** | 0.323 | 0.135 | 0.162 | -0.524 | -0.159 | -0.176 | -0.017 | -0.698 | 0.173 | -0.007 | 0.039 |
| **Hand Wing Index** | 0.105 | -0.740 | -0.001 | -0.028 | -0.099 | -0.270 | 0.104 | -0.002 | -0.155 | 0.344 | 0.452 |
| **Secondary1** | 0.338 | 0.148 | -0.303 | 0.023 | 0.055 | 0.351 | 0.562 | -0.020 | 0.044 | 0.553 | -0.147 |
| **Kipps Distance** | 0.269 | -0.520 | -0.108 | 0.044 | -0.126 | -0.037 | -0.091 | 0.021 | 0.222 | -0.169 | -0.734 |
| **Tail Length** | 0.296 | 0.042 | -0.345 | -0.084 | 0.779 | -0.282 | -0.302 | 0.012 | -0.068 | 0.040 | 0.013 |

**Supplementary Table 2** Tukey Honest Significant Differences of ANOVA describing variation in normalised root mean square error (normalised by standard deviation) as a factor of the percentage of species missing data, the metric, and whether data removal was random or biased. As the three-way interaction between metric, completeness and bias scenario was important for describing variation in normalised root mean square error, values are only given for comparisons including all three variables. For simplicity, only like-for-like comparisons that change one variable (changing variable in bold) are shown.

| Comparison | Difference (NRMSE, % SD) | p-value |
|---|---:|---:|
| **70%**: FUSE: Random-**30%**: FUSE: Random | 24.432 | <0.001 |
| 30%: **PRUNE**: Random-30%: **FUSE**: Random | -14.894 | <0.001 |
| 30%: FUSE: **Biased**-30%: FUSE: **Random** | -7.494 | <0.001 |
| 70%: **PRUNE**: Random-70%: **FUSE**: Random | -5.66 | <0.001 |
| 70%: FUSE: **Biased**-70%: FUSE: **Random** | -2.803 | <0.001 |
| **70%**: PRUNE: Random-**30%**: PRUNE: Random | 33.666 | <0.001 |
| 30%: PRUNE: **Biased**-30%: PRUNE: **Random** | -5.144 | <0.001 |
| 70%: PRUNE: **Biased**-70%: PRUNE: **Random** | 5.846 | <0.001 |
| **70%**: FUSE: Biased-**30%**: FUSE: Biased | 29.123 | <0.001 |
| 30%: **PRUNE**: Biased-30%: **FUSE**: Biased | -12.544 | <0.001 |
| **70%**: PRUNE: Biased-**70%**: FUSE: Biased | 2.989 | <0.001 |
| **70%**: PRUNE: Biased-**30%**: PRUNE: Biased | 44.656 | <0.001 |

### Supplementary Figures

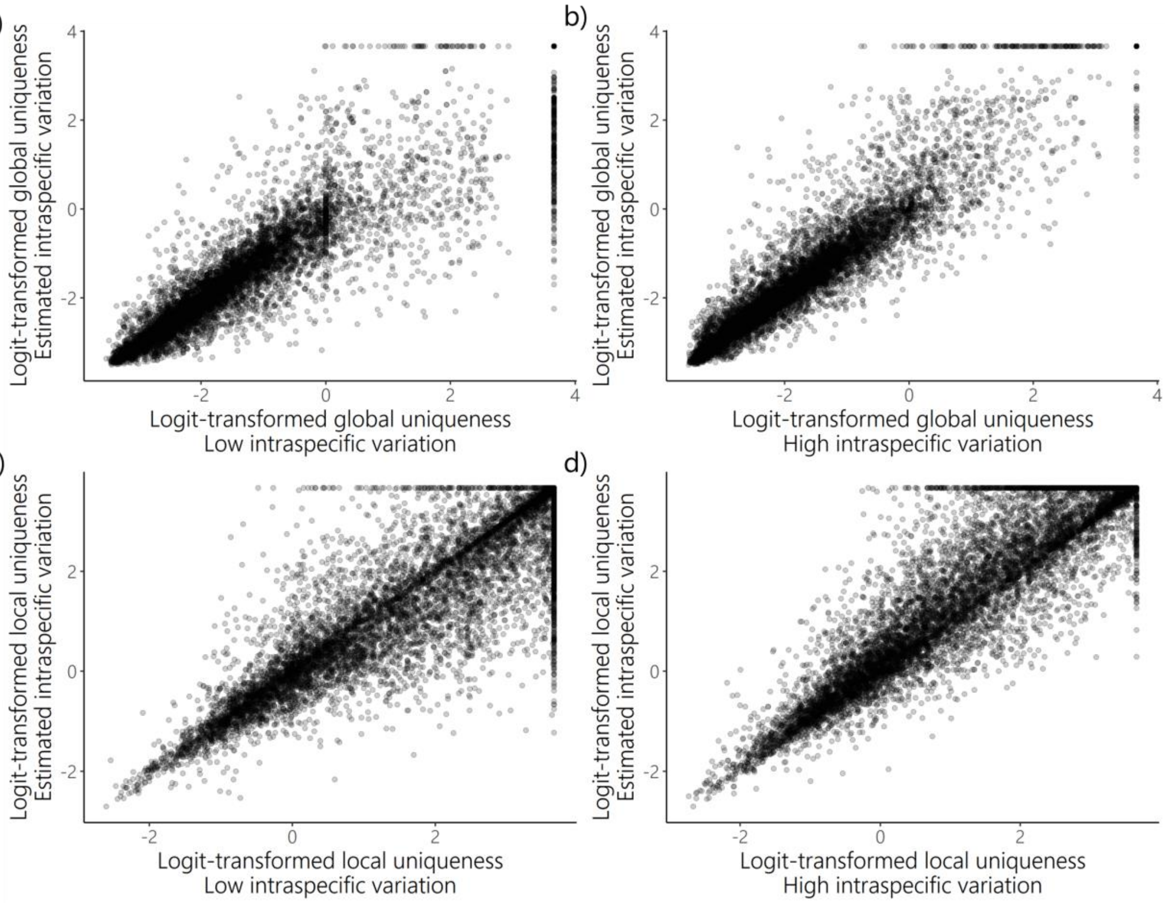


**Supplementary Figure 1** Logit-transformed global uniqueness (a and b, n=11 038 species) and local uniqueness (c and d, n= 10 859 species) where species probability distributions were constructed using standard deviation estimated with a bandwidth selector (estimated intraspecific variation, y axis in all panels), where standard deviation was decreased by 20% relative to that estimated with a bandwidth selector (low intraspecific variation, x axis in panels a and c), and where standard deviation was increased by 20% relative to that estimated with a bandwidth selector (high intraspecific variation, x axis in panels b and d).

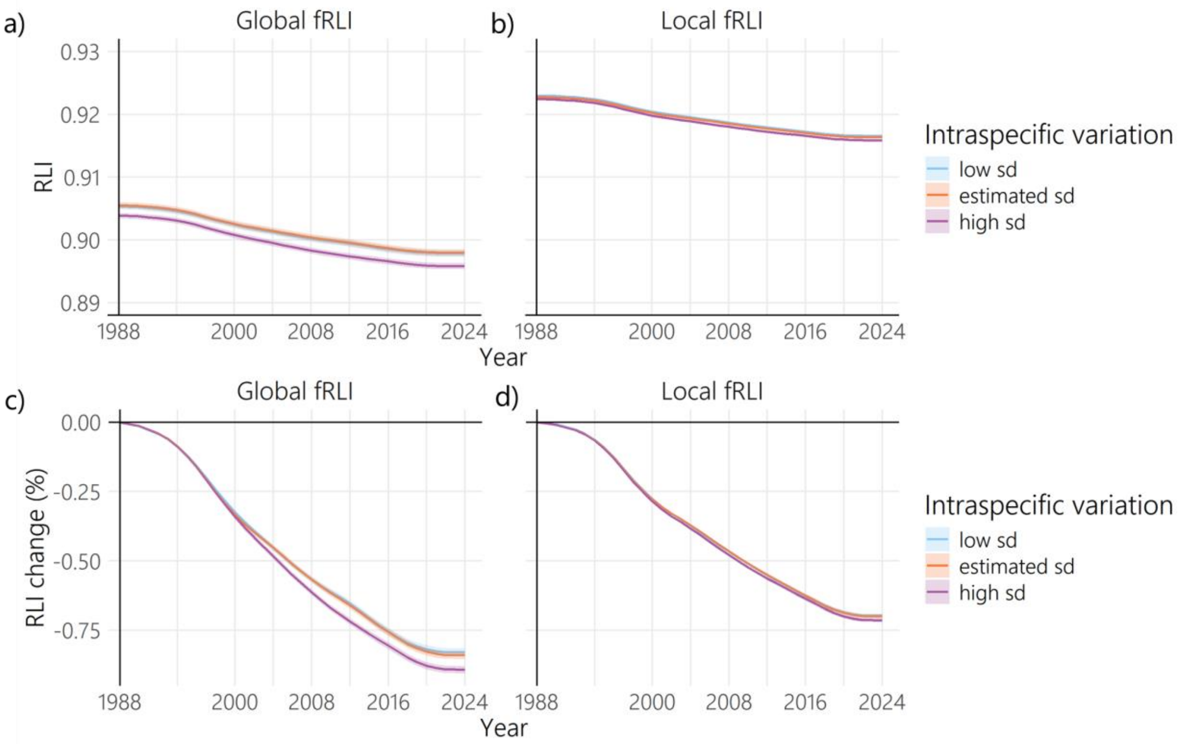


**Supplementary Figure 2** a) Global functional Red List Index (fRLI), b) local fRLI, c) global fRLI change between 1988 and 2024, and d) local fRLI change between 1988 and 2024, where species trait probability distributions were constructed using standard deviation estimated with a bandwidth selector (estimated sd), where standard deviation was decreased by 20% (relative to that estimated with a bandwidth selector, low sd), and where standard deviation was increased by 20% relative to that estimated with a bandwidth selector (high sd). fRLI calculations included 11 044 extant and recently extinct species. Error bands (shading) around the lines represent mean ± 2 x standard deviation of 1000 simulations where 40 Data Deficient species were assigned extinction risk trajectories at random from species with extinction risk data (of which three species were also missing range data), six species with missing trait data were assigned global functional uniqueness at random from global functional uniqueness of species with trait data, and 179 species with missing range data and six species with missing trait data were assigned local uniqueness at random from local uniqueness of species with trait and range data. A five-year moving average was calculated as described by Butchart et al. (2010) and back-casting was carried out as described in Butchart et al. (2007).

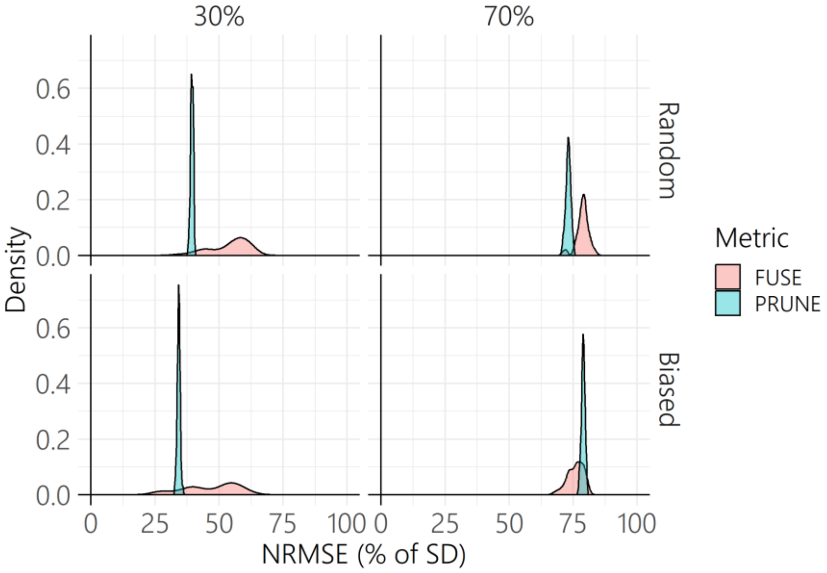


**Supplementary Figure 3** Density distributions of the root mean square error normalised by standard deviation (NRMSE (% of SD)) of functional uniqueness estimates for incomplete datasets, with 30% and 70% of species trait data missing and with random and biased data removal. Two functional uniqueness metrics were compared, the uniqueness metric based on nearest neighbours which underlies the FUSE (functionally unique, specialised and endangered) metric (Pimento et al., 2023) and probabilistic uniqueness calculated using trait probability densities (PRUNE).

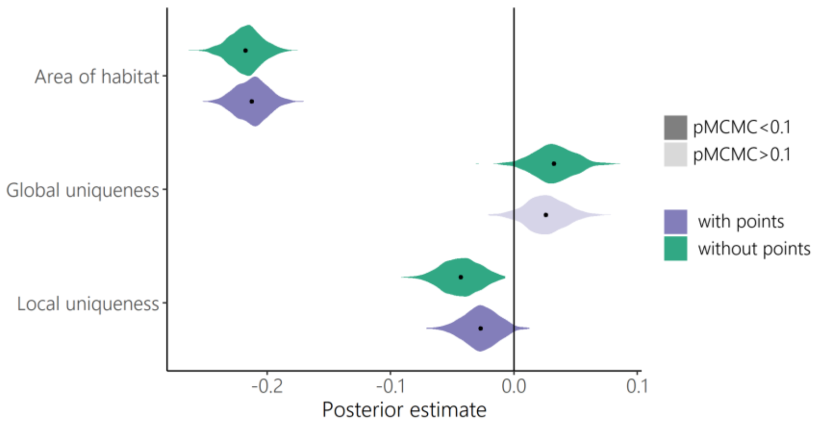


**Supplementary Figure 4** Posterior values from multi-response Monte-Carlo generalized linear mixed models showing the relationships between area of habitat, global uniqueness and mean local uniqueness and a) protected area coverage for resident areas of species' habitats including buffered centroids of area-based conservation sites where area was reported (as in the main text, with points) and b) excluding area-based conservation sites that were reported as points (without points). Global uniqueness and local uniqueness values were logit transformed, the mean logit-transformed local uniqueness value across species' area of habitat was calculated, then logit-transformed global uniqueness and mean logit-transformed local uniqueness were centred and scaled to allow for comparison of effect size between variables. Area of habitat was log10 transformed and then centred and scaled. We sampled 1000 posterior estimates (grey points show individual estimates), 8687 species were included in both models. pMCMC=0.1 was the threshold for significance.

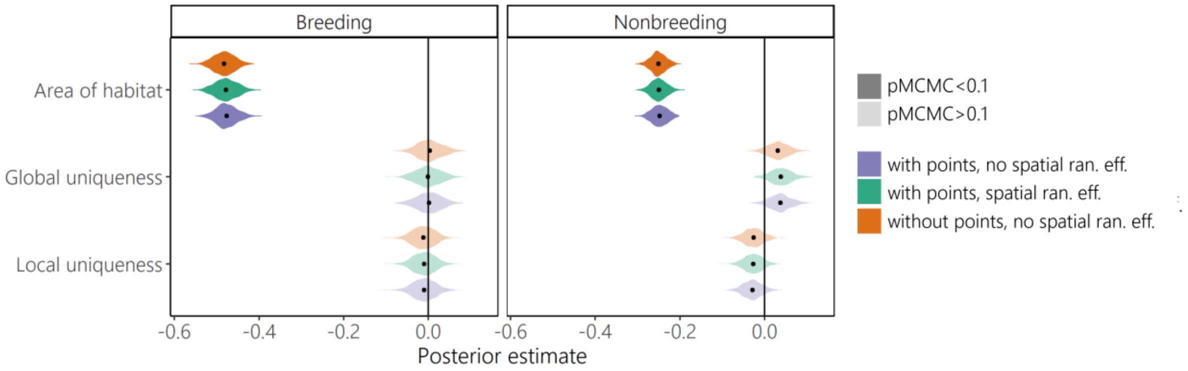


**Supplementary Figure 5** Posterior values from a multi-response Monte-Carlo generalized linear mixed models showing the relationships between area of habitat, global uniqueness and mean local uniqueness and protected area coverage for breeding areas of migratory species' habitats (n=2297 species) and non-breeding area of migratory species' habitats (n=2201 species), where points from the WDPA and WDCOEM were included as buffered centroids (with points) and where points were excluded. Models were run with three spatial eigenvectors as random effects (spatial ran. eff.), and without spatial random effects (no spatial ran. eff.). pMCMC=0.1 was the threshold for significance. Global uniqueness and local uniqueness values were logit transformed, the mean logit-transformed local uniqueness value across species' area of habitat was calculated, then logit-transformed global uniqueness and mean logit-transformed local uniqueness were centred and scaled to allow for comparison of effect size between variables (variables were centred and scaled for resident, breeding and non-breeding models separately). Area of habitat was log10 transformed and then centred and scaled. Migratory species do not necessarily have both breeding and non-breeding areas of habitat, which is particularly the case for species which are partially migratory, and for some migratory species breeding or non-breeding areas of habitat are not known. For every model we sampled 1000 posterior estimates.

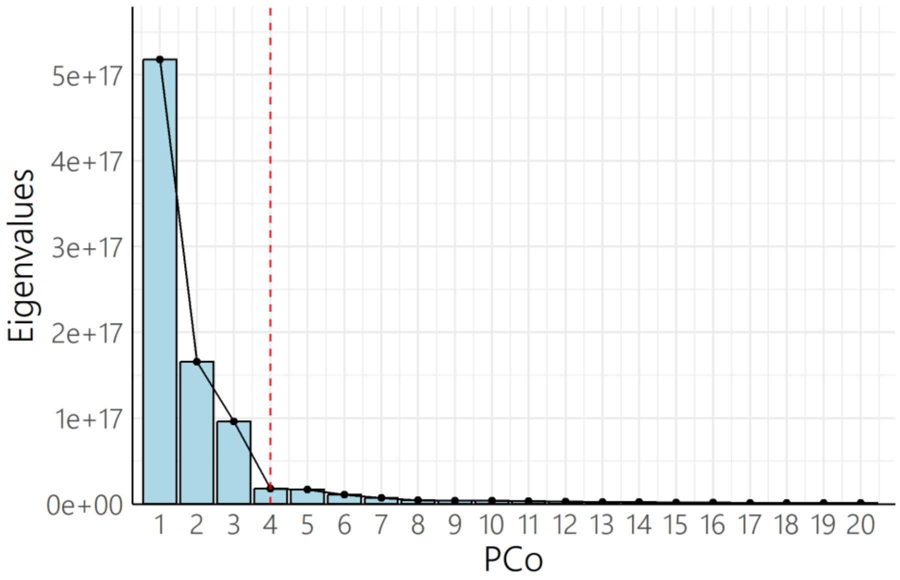


**Supplementary Figure 6** Eigenvalues for the first 20 principal coordinates axes (eigenvectors) of a Haversine distance matrix between species centroids (n=10,806 species). The dotted line shows the elbow of the scree plot after which adding more principal coordinate axes describes minimal additional variance.
